# Membrane Tethering of Honeybee Antimicrobial Peptides in *Drosophila* Enhances Pathogen Defense at the Cost of Stress-Induced Host Vulnerability

**DOI:** 10.1101/2025.07.17.665347

**Authors:** Yanan Wei, Yanying Sun, Xinyue Zhou, Doyoun Kim, Jihyeon Lee, Jeong Kyu Bang, Woo Jae Kim

**Author notes:** These authors contributed equally (Yanan Wei, Yanying Sun, Xinyue Zhou). YW YS XZ DK JL: JH JKB.

## Abstract

Antimicrobial peptides (AMPs) represent a promising alternative to conventional antibiotics in combating multidrug-resistant pathogens, yet their clinical translation is hindered by proteolytic instability, cytotoxicity, and poor bioavailability. Here, we demonstrate that glycosylphosphatidylinositol (GPI)-mediated membrane tethering of honeybee defensin1 (Def1) in *Drosophila melanogaster* enhances its antimicrobial efficacy by ∼100-fold compared to secreted or untethered forms, while preserving physiological and behavioral integrity under baseline conditions. Using a genetically engineered *Drosophila* model, we expressed three Def1 variants: native (Def1), secreted (s-Def1), and membrane-tethered (t-Def1). Flies expressing t-Def1 exhibited superior bacterial clearance of *Pseudomonas aeruginosa* and improved survival post-infection, with no adverse effects on locomotion, courtship, or sleep architecture. However, under stress paradigms—including sleep deprivation and dextran sulfate sodium (DSS)-induced gut injury—t-Def1 exacerbated intestinal barrier dysfunction, as evidenced by elevated Smurf phenotype incidence, highlighting a trade-off between antimicrobial potency and epithelial vulnerability. Our work establishes *Drosophila* as a powerful platform for dissecting AMP mechanisms and engineering spatially targeted therapies, offering translational insights for pollinator health and human infectious disease management. These results advocate for iterative refinement of membrane-anchoring strategies to balance therapeutic efficacy with host safety, advancing the development of next-generation AMPs with minimized off-target effects.

## INTRODUCTION

Antimicrobial peptides (AMPs) are evolutionarily conserved components of innate immunity, serving as first-line defenders against pathogens through mechanisms ranging from membrane disruption to intracellular targeting of essential bacterial processes^[1]^. These short cationic peptides exhibit broad-spectrum activity against bacteria, fungi, viruses, and parasites, making them promising alternatives to conventional antibiotics amid rising antimicrobial resistance^[2]^. The global rise of multidrug-resistant pathogens has reignited pharmaceutical interest in AMPs as potential therapeutics, with over 60 candidates in clinical trials for infections, wound healing, and immunomodulation^[3]^. For instance, LL-37, a human cathelicidin, is in Phase II trials for diabetic foot ulcers^[4]^, while synthetic AMPs like pexiganan target antibiotic-resistant skin infections^[5]^. Despite this progress, challenges such as proteolytic degradation, high production costs, and cytotoxicity limit widespread clinical adoption. The global AMP market, valued at $3.8 billion in 2022, is projected to grow at 6.5% annually, driven by demand for novel antimicrobial strategies^[6]^. Understanding AMP mechanisms and optimizing their efficacy thus remains a pressing biomedical priority.

Honeybees (*Apis mellifera*), vital pollinators facing population declines linked to pathogens like *Paenibacillus larvae*, rely heavily on AMPs such as defensin1, abaecin, and hymenoptaecin for immune defense^[7]^. Forager bees exhibit upregulated AMP expression in hypopharyngeal and mandibular glands, suggesting roles in both immunity and food preservation^[8]^. However, the molecular mechanisms underlying their activity remain poorly characterized due to limited genetic tools in apian models^[9]^. Functional studies in bees are constrained by their complex social behavior, long life cycles, and ethical considerations, hindering mechanistic exploration^[10]^. To overcome these limitations, we leverage *Drosophila melanogaster*, a genetically tractable model organism with conserved immune pathways, including the Toll and Imd signaling cascades that regulate AMP production^[11]^. The fruit fly’s unparalleled genetic toolkit—including RNAi, CRISPR-Cas9, and tissue-specific overexpression systems—enables precise dissection of AMP function *in vivo*, offering a robust platform to study heterologous honeybee AMPs in a controlled, high-throughput manner^[12,13]^.

Both honeybees and fruit flies rely on the Toll and Imd innate immune signaling pathways to regulate the expression of antimicrobial peptides (AMPs). Their core immune components (e.g., Toll receptors, NF-κB transcription factors) and regulatory networks exhibit remarkable evolutionary conservation^[14,15]^. Although honeybee Def1 and *Drosophila* AMPs (such as Drosomycin and Diptericin) show significant sequence divergence (<30% homology), they all maintain the conserved cysteine-stabilized αβ (CSαβ) structural fold^[16]^, demonstrating functional conservation at the structural level. This characteristic “pathway conservation with sequence divergence” makes *Drosophila* an excellent model system for studying heterologous AMPs (e.g., honeybee Def1), while avoiding functional interference with endogenous AMPs^[14]^.

Recent advances suggest that AMP activity can be enhanced by modifying their localization via membrane tethering^[17]^. Conventional membrane tethering strategies typically employ covalent lipid modifications (e.g., N-terminal myristoylation or palmitoylation) or transmembrane domains to anchor peptides. Myristoylation attaches a 14-carbon saturated fatty acid via an amide bond to glycine residues, while palmitoylation links 16-carbon chains through thioester bonds to cysteine residues^[18]^. Transmembrane domains use hydrophobic α-helices spanning the lipid bilayer. While membrane tethering represents a promising strategy to amplify the antimicrobial efficacy of AMPs, engineering such structural modifications *in vitro* remains technically demanding, as cell-free systems often fail to recapitulate the complex lipid interactions and spatial organization required for stable peptide anchoring. Given the technical barriers to achieving functional AMPs tethering *in vitro*, we engineered a genetically encoded system in which AMPs are covalently anchored to the extracellular leaflet of the plasma membrane via a glycosylphosphatidylinositol (GPI) modification. This approach bypasses the limitations of cell-free environments by leveraging the host’s endogenous machinery for precise spatial localization and stable membrane integration^[19–21]^. This *in vivo* platform ensures proper peptide orientation and lipid interactions, overcoming the instability inherent to synthetic membrane models^[22,23]^.

In this study, we demonstrate that GPI-tethering of honeybee defensin1 (Def1) in *Drosophila* dramatically amplifies its antimicrobial activity compared to its secreted form. Using transgenic flies expressing tethered defensin1 (t-Def1), we show that membrane anchoring enhances peptide stability and proximity to invading microbes, providing a mechanistic explanation for its superior performance. This approach not only clarifies the functional relevance of honeybee AMPs but also establishes a versatile paradigm for engineering AMPs with improved therapeutic potential. By bridging honeybee immunology with *Drosophila* genetics, our work advances understanding of AMP mechanisms and potential applications in pathogen defense. This hybrid model provides fundamental insights into honeybee AMP mechanisms while establishing a platform for developing antimicrobial strategies with particular relevance to pollinator health.

## RESULTS

### GPI-mediated Membrane Anchoring of Honeybee Defensin1 in *Drosophila* Reveals Spatial Localization Enhances Antimicrobial Efficacy

Defensin1 (Def1), a key honeybee AMP with broad-spectrum antimicrobial properties but poorly characterized mechanisms, was chosen to evaluate GPI tethering in *Drosophila*—a model enabling precise genetic manipulation to resolve how spatial anchoring optimizes peptide function *in vivo*^[24,25]^. By testing GPI-tethered Def1 in *Drosophila*, we leverage the fruit fly’s genetic tractability to dissect how membrane localization enhances peptide efficacy, bypassing the technical limitations of honeybee systems while providing insights applicable to both pollinator health and therapeutic AMP engineering.

To evaluate the antimicrobial activity of Def1, we engineered three transgenic strains expressing distinct defensin1 variants (Fig. 1A-B). The first, designated *upstream activation sequence (UAS)-Def1*, encodes the native defensin1 sequence, including its endogenous signal peptide and propeptide. The second, *UAS-s-Def1* (secreted defensin1), incorporates the α-bungarotoxin (α-Bgtx) signal peptide followed by the defensin1 sequence and a C-terminal hydrophilic linker (GGNEQKLISEEDLGN) to enhance extracellular secretion. The third variant, *UAS-t-Def1* (tethered defensin1), was designed by appending a glycosylphosphatidylino sitol (GPI) anchor sequence to the C-terminus of *UAS-s-Def1* to facilitate membrane tethering. These constructs were generated to systematically assess how subcellular localization and secretion mechanisms influence Def1’s functional efficacy (Fig. 1A-B)^[21,22]^. Bungarotoxin binds irreversibly to nicotinic acetylcholine receptors (nAChRs) via specific interactions with receptor subunits, blocking synaptic transmission^[26]^, while GPI (glycosylphosphatidylinositol) anchors proteins to the outer leaflet of the plasma membrane through a covalent linkage between the protein’s C-terminus and a preassembled glycolipid, enabling stable membrane attachment without transmembrane domains^[27]^. The GPI anchor is covalently linked to the C-terminus of t-Def1 through a transamidation reaction catalyzed by GPI transamidase, which cleaves the precursor’s C-terminal signal peptide and attaches a preassembled GPI glycolipid via an amide bond between the terminal ethanolamine moiety and Def1’s carboxyl group^[27]^. This modification embeds the peptide stably into the membrane outer leaflet without transmembrane domains.

**Fig. 1:**
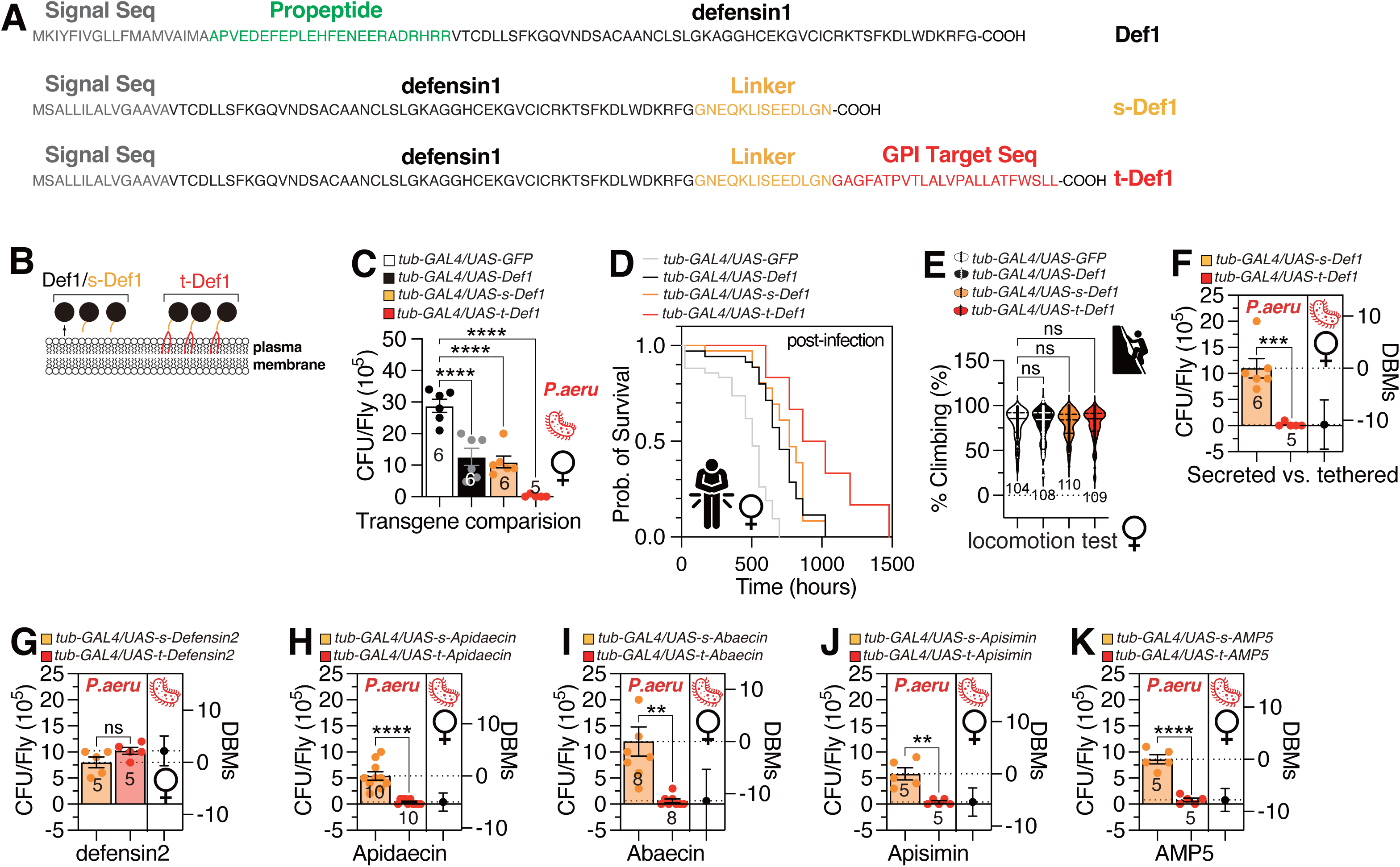
The diagram of honeybee Defensin1 related peptides and the antimicrobial effect of flies expressing honeybee AMPs. **A,** The amino acid sequences of native Defensin1(Def1), secretion format Defensin1 (s-Def1) and tethered form Defensin1(t-Def1), including signal sequences (blue), propeptide sequence (green), Defensin1 chain sequence (black), linker sequence (yellow), GPI target sequence (red). **B,** The diagram of Def1, s-Def1, t-Def1, and their positions to cell membrane. **C,** Pathogen load of female flies expressing Def1, s-Def1, and t-Def1 compared with control (GFP expressing flies) by *tub-GAL4* following a 12-hour oral feeding of *P. aeruginosa* culture. CFU stands for Colony-Forming Unit. **D,** Survival curves of female flies expressing Def1, s-Def1, and t-Def1 compared with control (GFP expressing flies) by *tub-GAL4* after infection by *P. aeruginosa* culture. **E,** The climbing ability of female flies expressing Def1, s-Def1, and t-Def1 compared with control (GFP expressing flies) by *tub-GAL4*, numbers shown are sample size of each condition. **F-K,** Pathogen load of female flies expressing **F,** s-Defensin1 and t-Defensin1, **G,** s-Defensin2 and t-Defensin2, **H,** s-Apidaecin and t-Apidaecin, **I,** s-Abaecin and t-Abaecin, **J,** s-Apisimin and t-Apisimin, **K,** s-AMP5 and t-AMP5. DBMs represent difference between means, which is a statistical measure that quantifies the average discrepancy between two groups. Numbers shown are sample sizes for each condition. The bar plots with dots represent the colony forming unit (CFU) per fly. The numerical values beneath the bar plots indicate the count of flies that were successfully tested. The mean value and standard error are labeled within the bar plot. DBMs represent the ‘difference between means’ for the evaluation of estimation statistics (See **METHODS**). Asterisks represent significant differences, as revealed by the unpaired Student’s t test, and ns represents non-significant differences (*\*p<0.05, **p<0.01, ***p< 0.001, ****p< 0.0001*). The identical analytical approach employed for the two sample comparision is maintained for the subsequent data presented.

To compare the AMP activity of Def1 variants, we measured pathogen load in female flies expressing each form after a 12-hour oral infection with *Pseudomonas aeruginosa* (Fig. 1C). Flies expressing Def1, s-Def1 (secreted), or t-Def1 (membrane-tethered) under *tub-GAL4* exhibited significantly reduced bacterial burdens. Notably, t-Def1 demonstrated ∼100-fold greater antimicrobial efficacy than Def1 or s-Def1, as shown by bacterial clearance (Fig. 1C) and post-infection survival (Fig. 1D). This enhanced activity was independent of mRNA levels, as all variants showed comparable expression (Fig. S1A). Locomotion in females (Fig. 1E) and males (Fig. S1B) was unaffected by Def1 expression. Similarly, male courtship behavior (courtship index, copulation latency) remained unchanged across variants (Fig. S1C-D), indicating no behavioral impacts of Def1 expression.

To investigate the influence of membrane tethering on the functional efficacy of honeybee AMPs, multiple AMP variants were engineered in both secreted and membrane-anchored configurations. A comparative analysis of their antimicrobial potency revealed that membrane-tethered AMPs generally exhibited enhanced activity relative to their secreted counterparts. However, defensin2 (Def2) represented an exception, as both its secreted and tethered forms demonstrated equivalent antimicrobial effects (Fig. 1F–K). These findings indicate that the incorporation of a flexible polypeptide linker combined with a GPI-anchoring sequence substantially potentiates the antimicrobial function of honeybee AMPs when anchored to cellular membranes.

While initial characterization demonstrated that membrane-tethered t-Def1 provided potent protection against *Pseudomonas aeruginosa* infection (Fig. 1F), we further investigated whether this engineered AMP retained the broad-spectrum activity characteristic of natural antimicrobial peptides. To directly test efficacy against Gram-positive pathogens, we challenged t-Def1-expressing flies with *Enterococcus faecalis*. Strikingly, t-Def1 expression conferred robust protection against this Gram-positive bacterium (Fig. S1E), with survival rates significantly exceeding controls. This expanded pathogen profile confirms that t-Def1’s enhanced antimicrobial activity is not limited to Gram-negative organisms but extends across bacterial classifications, consistent with the inherent broad-spectrum functionality of defensins. The conserved efficacy against structurally distinct pathogens reinforces that t-Def1’s engineered localization mechanistically amplifies its innate antimicrobial properties.

To rigorously attribute observed phenotypes to the engineered defensin constructs rather than genetic background or insertion site effects, we expanded our control strategy beyond GFP-expressing flies. We introduced two critical additional control groups: (1) wild-type Canton-S (CS) flies, and (2) transgenic flies carrying an empty UAS vector integrated at the identical genomic attP40 site used for our AMP constructs. Comparative analysis of pathogen load across these control groups revealed no significant differences between wild-type (CS) and empty UAS vector *(+/attP40*) (Fig. S1F). This equivalence confirms that neither the attP40 insertion site nor the general genetic manipulation background contributes measurably to the phenotypic outcomes. Consequently, the pronounced reductions in pathogen load and associated physiological phenotypes observed in t-Def1 and t-Def2 expressing lines (Figs. 1F-K) are unequivocally attributable to the specific activity of the engineered AMP constructs.

To address the critical need for direct validation of protein expression levels and localization for our tethered defensin constructs (t-Def1, t-Def2), we performed targeted experiments beyond initial mRNA quantification. To directly assess expression levels of tethered defensins (t-Def1/t-Def2), we performed c-myc-targeted ELISA despite lacking defensin-specific antibodies. Quantitative analysis confirmed comparable protein expression between t-Def1 and t-Def2 (Fig. S1G-H; ns), consistent with their equivalent mRNA levels (Fig. S1A). This equivalence stands in stark contrast to t-Def1’s dramatic ∼100-fold superior antimicrobial efficacy against *P. aeruginosa* (Fig. 1C), demonstrating that functional differences arise from post-translational mechanisms rather than expression quantity. We next resolved the cellular localization of tethered defensins using c-myc immunofluorescence in enterocytes. Despite technical challenges from membrane-proximal epitope burial, we detected distinct c-myc signal specifically in t-Def1-expressing guts, with pronounced plasma membrane enrichment (Fig. S1I-J). This pattern aligns with t-Def1’s GPI-anchor design and provides direct structural evidence for successful membrane tethering. Functionally, this membrane localization correlated with enhanced pathogen clearance: When challenged with GFP-expressing *P. aeruginosa*, t-Def1 guts exhibited significantly reduced bacterial colonization compared to genetic controls (Fig. S1J). This spatial correlation between membrane-localized defensin and bacterial clearance provides mechanistic validation that tethering positions AMPs at critical pathogen-entry interfaces to maximize antimicrobial impact.

Notably, t-Def2 showed no enhanced antimicrobial activity compared to s-Def2 (Fig. 1G). AlphaFold structural predictions revealed that t-Def1 maintained the native cysteine-stabilized αβ (CSαβ) fold (predicted Local Distance Difference Test > 90), while t-Def2 exhibited low-confidence (pLDDT < 50) at the linker-GPI junction (Fig. S2A-B), suggesting that structural destabilization may underlie its functional insensitivity to membrane tethering. This divergence highlights the importance of structural integrity for GPI-anchored AMP efficacy.

### Membrane-tethered Defensin1 Enhances Antimicrobial Activity without Disrupting Locomotor or Sleep Behavior in *Drosophila*

The membrane-tethered defensin1 (t-Def1) variant exhibited significantly enhanced antimicrobial activity compared to its secreted counterpart (s-Def1), while no adverse effects on climbing ability or male courtship behavior were observed (Fig. 1 and S1). To further assess potential physiological or behavioral impacts of t-Def1 expression, horizontal locomotion was analyzed in both isolated and grouped flies. Flies expressing t-Def1 displayed modestly reduced horizontal movement velocity relative to s-Def1-expressing individuals (Fig. 2A). Although stop frequency, rotation events, and spatial distribution patterns remained comparable between groups (Fig. 2B–D), t-Def1 flies exhibited shorter total trajectory lengths, likely attributable to reduced mobility (Fig. 2E–F). Group behavioral assays further demonstrated equivalent inter-fly proximity, wall distance, angular velocity, and vertical speed between t-Def1 and s-Def1 flies (Fig. 2G–J and S3A). However, t-Def1 flies maintained slower horizontal speeds in collective settings (Fig. 2K), consistent with individual locomotion data (Fig. 2A).

**Fig. 2:**
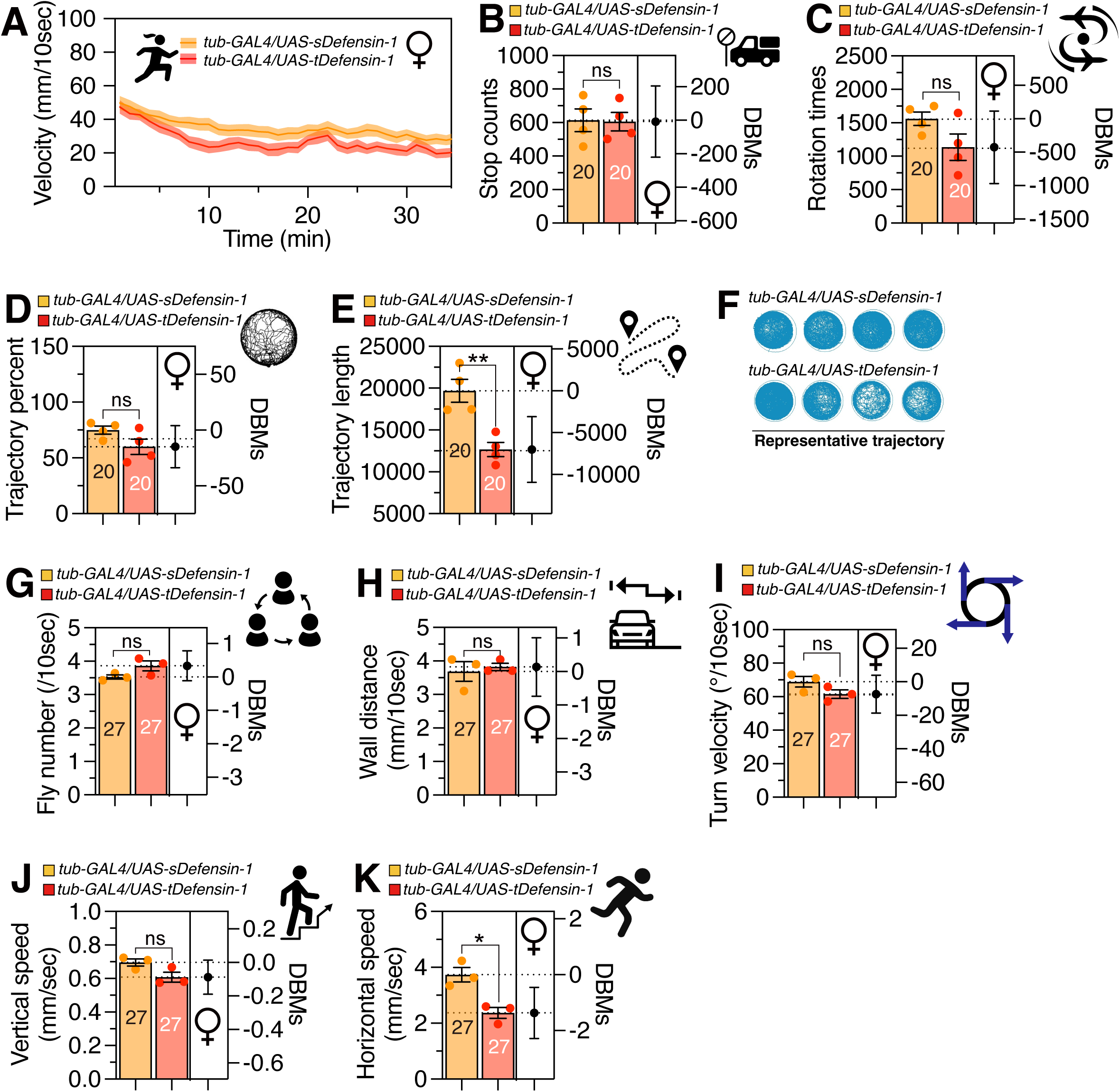
Locomotion ability of female flies expressing s-Def1 and t-Def1. The **A,** velocity, **B,** stop counts, **C,** rotation times, **D,** trajectory percent, and **E,** trajectory length of single female s-Def1 and t-Def1 flies. For rotation time, it is defined as the counts of flies turning over 45° in 2 seconds. **F,** The representative trajectory recording graph. The **G,** fly number, **H,** wall distance, **I,** turn velocity, **J,** vertical speed and **K,** horizontal speed of grouped female s-Def1 and t-Def1 flies. Numbers shown are sample sizes for each condition. For the fly number, it refers to the number of flies each single fly in the group encountered in a diameter of 2mm. And horizontal as well as vertical speed, is defined as average speed of flies moving in horizontal and vertical directions respectively.

To assess the potential effects of membrane-tethered defensin1 (t-Def1) on sleep regulation, sleep architecture was analyzed in flies with ubiquitous expression of either secreted (s-Def1) or tethered t-Def1 under the *tub-GAL4* driver (Fig. 3). Sleep profiles, quantified as the percentage of time spent inactive in 30-minute intervals across a 24-hour light-dark cycle, revealed closely matched patterns between s-Def1– and t-Def1-expressing flies (Fig. 3A). Total sleep duration (Fig. 3B), daytime sleep duration (Fig. 3C), nighttime sleep duration (Fig. 3D), total sleep episodes (Fig. 3E), daytime sleep episodes (Fig. 3F), nighttime sleep episodes (Fig. 3G), and sleep latency (Fig. S4A–B) were statistically indistinguishable between s-Def1– and t-Def-expressing flies. These findings collectively indicate that constitutive expression of t-Def1 expression does not overtly impair key physiological or behavioral parameters under baseline conditions, though subtle reductions in locomotor velocity and trajectory length were noted.

**Fig. 3:**
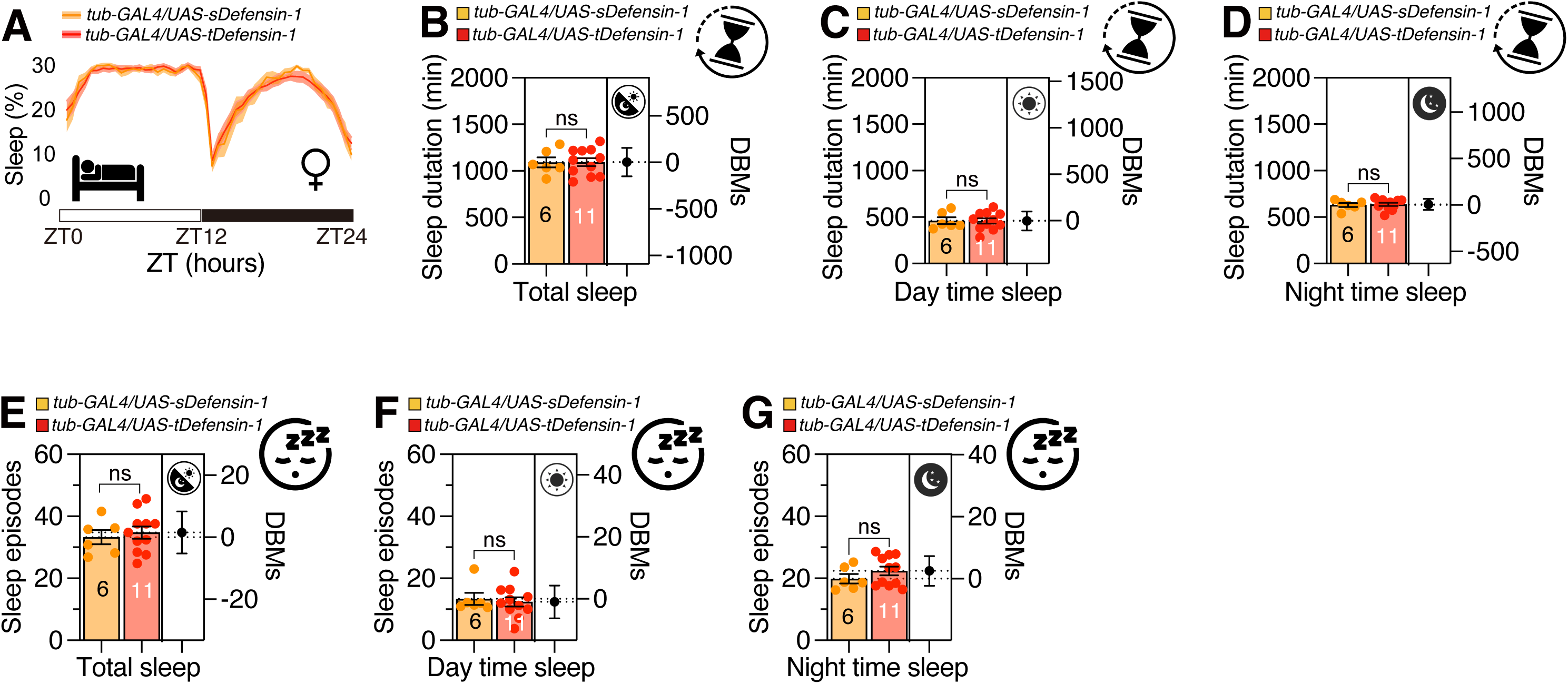
The sleep pattern of flies with ubiquitous expression of s-Def1 and t-Def1. **A**, Average sleep profiles (proportion of time spent sleeping in consecutive 30-min segments during a 24-h light-dark (LD) cycle) of female flies ubiquitously expressing s-Def1 and t-Def1. **B-D,** Quantification of sleep duration for s-Def1 and t-Def1 expression flies by *tub-GAL4* driver. Data are presented as box plots with individual data points. **E-G**, Quantification of sleep episodes for s-Def1 and t-Def1 expression flies. Data are presented as box plots with individual data points. (n=6, n=11 respectively).

### Membrane-tethered Defensin1 Exacerbates Stress-Induced Epithelial Compromise in *Drosophila*

While AMPs provide robust defense against pathogens, their cytotoxicity toward host cells becomes apparent under conditions where phosphatidylserine (PS)—a negatively charged phospholipid—is exposed on the outer membrane leaflet during stress^[28,29]^. However, mechanisms regulating AMP cytotoxicity in diverse cellular contexts remain underexplored. To evaluate the capacity of the engineered peptide t-Def1 to counteract host cell damage mediated by AMPs, we assessed its effects under both physiological and stress-induced conditions.

To evaluate intestinal barrier integrity in response to t-Def1 expression, we employed the Smurf assay, a non-invasive method for quantifying gut permeability in *Drosophila melanogaster*. This assay detects systemic leakage of the non-absorbable dye FD&C Blue No. 1 into the hemolymph, a phenomenon indicative of disrupted epithelial tight junctions^[30,31]^. The Smurf phenotype—characterized by whole-body blue coloration—serves as a validated biomarker of intestinal dysfunction, correlating strongly with physiological decline and mortality in aging flies^[32]^ and various stresses^[33,34]^.

Under non-stressed conditions, no significant difference in Smurf phenotype intensity—a marker of intestinal permeability—was observed between s-Def1– and t-Def1-expressing females (Fig. 4A). Following 6 hours or 24 hours of sleep deprivation (SD) and 6 hours of blue dye feeding, however, t-Def1 flies exhibited elevated abdominal dye leakage compared to s-Def1 controls (Fig. 4B, S5A). To further probe stress susceptibility, flies were exposed to dextran sulfate sodium (DSS), a chemical agent known to induce mucosal injury and compromise gut barrier function in *Drosophila* (Fig. 4C)^[35,36]^. After treatment with 3% DSS, t-Def1-expressing flies demonstrated a significantly higher incidence of the Smurf phenotype relative to s-Def1 flies (Fig. 4C). These results collectively indicate that membrane tethering of Def1 amplifies epithelial damage under both physiological and chemical stress paradigms.

**Fig. 4:**
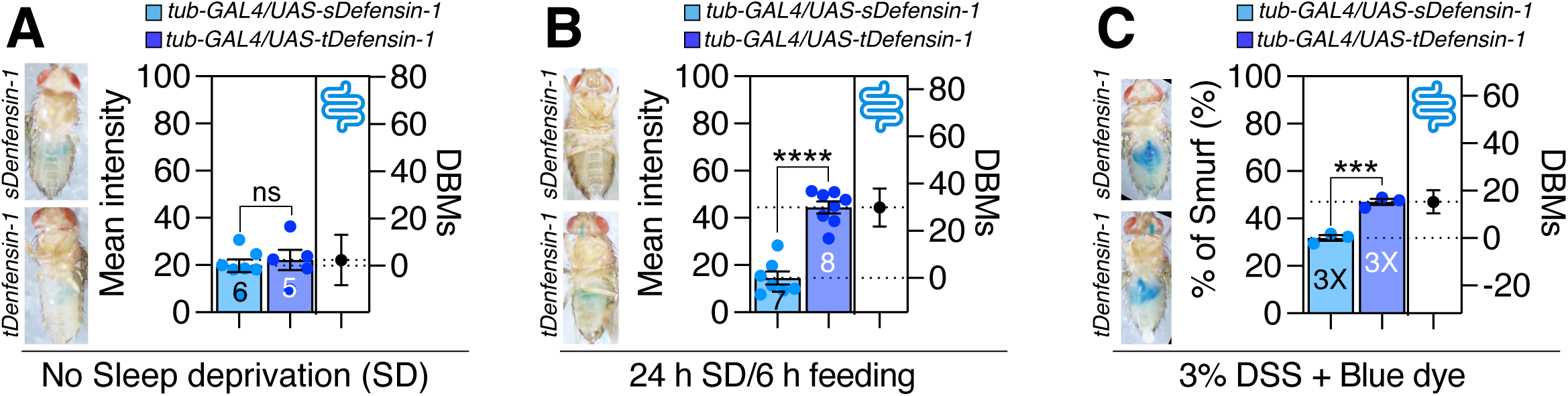
Smurf results of female s-Def1 and t-Def1 flies in normal condition and induced by different types of stress. **A,** The mean intensity of Smurf assay flies expressing s-Def1 compared with t-Def1 under normal condition. Numbers shown are sample sizes for each condition, and one representative figure was shown of each condition. **B,** The mean intensity of Smurf assay flies expressing s-Def1 compared with t-Def1 after 24 hours sleep deprivation and 6 hours dye food feeding. **C,** The percentage of flies (n=11 and 12 respectively, repeated for 3 times) showing Smurf phenotype after a 3% DSS feeding with blue dye. The experiment was repeated three times and one representative figure was shown for each condition.

## DISCUSSION

Our findings demonstrate that GPI-mediated membrane tethering of honeybee Def1 in *Drosophila melanogaster* significantly amplifies its antimicrobial efficacy while preserving key physiological and behavioral functions under baseline conditions. Notably, wild-type *Drosophila* employs endogenous AMPs such as Drosomycin (antifungal) and Diptericin (antibacterial), which are regulated by the Toll and Imd pathways, respectively. While these peptides share the cysteine-stabilized αβ (CSαβ) fold with honeybee Def1, their low sequence homology (<30%) minimizes functional overlap^[16,37]^. This divergence allowed t-Def1 to operate without interfering with native immune responses, as evidenced by unaltered endogenous AMP expression in our model. By engineering three distinct variants— native Def1, secreted s-Def1, and membrane-tethered t-Def1—we established that spatial anchoring via GPI modification enhances bacterial clearance by ∼100-fold compared to secreted or untethered forms, as evidenced by reduced *Pseudomonas aeruginosa* pathogen load and improved post-infection survival in transgenic flies. This enhanced activity was independent of transcriptional regulation, as mRNA levels remained comparable across variants, and critically, t-Def1 expression did not impair locomotion, courtship behavior, or sleep architecture, underscoring its biocompatibility in unstressed organisms. However, under stress paradigms, such as sleep deprivation or chemical challenge with DSS, t-Def1 exacerbated intestinal barrier dysfunction, as revealed by elevated Smurf phenotype incidence and dye leakage, suggesting a context-dependent trade-off between antimicrobial potency and epithelial integrity. The use of *Drosophila* as a genetically tractable model enabled precise dissection of these dual roles, bridging honeybee AMP biology to conserved mechanisms of immune defense and stress response. These results highlight the potential of membrane tethering as a strategy to optimize AMP activity while underscoring the need to balance therapeutic efficacy with host tissue vulnerability. The general applicability of GPI-anchoring is further substantiated by our independent validation of diverse AMP classes (e.g., cecropins, defensins) in *Drosophila* models, where structural optimization preserved immunomodulatory function^[38]^. This confirms the strategy’s compatibility beyond honeybee-derived peptides. By leveraging evolutionary conservation and *Drosophila*’s genetic toolkit, this work advances translational insights into AMP engineering, offering a roadmap for developing targeted antimicrobial therapies with minimized off-target effects, applicable to both pollinator conservation and human infectious disease management.

Insect defensins, including Apis mellifera Def1 and Def2, adopt the cysteine-stabilized αβ (CSαβ) motif ^[16]^, characterized by an N-terminal loop, an α-helix, and two antiparallel β-strands, stabilized by three conserved disulfide bonds. UniProt entries for Def1 (P17722 and Q5J8R1) indicate these bonds form between Cys46-Cys74, Cys60-Cys79, and Cys64-Cys81 in the precursor sequence, ensuring structural stability essential for antimicrobial activity.

The mechanism by which Def1 exerts its antimicrobial effects primarily involves disrupting the integrity of the bacterial cell membrane. It is proposed that defensins can achieve this by inserting themselves into the lipid bilayer of the bacterial membrane, leading to increased permeability and ultimately cell death^[16]^. Number of studies suggest that defensins can form channels within the plasma membrane of target cells. This channel formation can result in the leakage of essential cytoplasmic components, such as potassium ions, leading to a decrease in intracellular ATP levels and the inhibition of vital respiration processes^[39]^. More generally, the antimicrobial action of defensins can be attributed to either a direct disruption of the microbial cell membrane or an interference with essential metabolic processes within the microbial cell.

The AlphaFold prediction of both native Def-1 and Def-2 indicated that they adopt typical β-defensin folds with three disulfide bonds formed by conserved cysteine residues (Fig. S2A, B, D, and F). To address the functional discrepancy between GPI-tethered Def-1 (tDef-1) and tDef-2, the defensin structures incorporating linker and GPI-tethering sequences were modeled under identical conditions using AlphaFold.

The prediction confidence levels for the linker and GPI-tethered sequences in the modeled Def-1 were low, whereas the other regions demonstrated relatively high confidence (Fig. S2A). In contrast, the structural confidence for modeled Def-2 showed poor scores at the amino-terminal, linker, and GPI-tethered regions (Fig. S2B). Notably, the modeled sDef-2 and tDef-2 exhibited an anti-parallel structure in the linker region, resulting in significant amino-terminal structural alterations, while the structural integrity of Def-1 remained intact after modification (Fig. S2C and E). The substantial conformational change at the amino terminus of tDef-2 appears to impact the formation of the first cysteine pair’s disulfide bond, a crucial structural feature for the proper functional folding of β-defensin. Although AlphaFold’s lower confidence scores typically correlate with intrinsic disorder or high flexibility, for the GPI-anchored Def2, this uncertainty might, paradoxically, reflect a structural alteration from its expected wild-type structure, suggesting a functionally relevant structural alteration in these critical regions, especially N-terminal alpha helix. To generalize GPI-anchoring as a strategy for AMP optimization, future studies should refine linker flexibility and GPI-anchoring motifs to preserve the structural fidelity of diverse defensins, ensuring membrane localization enhances rather than compromises their antimicrobial mechanisms.

The lack of enhanced antimicrobial efficacy observed with GPI-tethered Def2 (t-Def2) may stem from structural perturbations induced by the fusion of its flexible linker and GPI-anchoring sequence. Insect defensins, including *Apis mellifera* Def1, rely on a conserved cysteine-stabilized αβ (CSαβ) motif—a compact tertiary architecture maintained by three disulfide bonds critical for membrane interaction and antimicrobial activity. AlphaFold structural predictions reveal that the addition of the linker and GPI-target sequence disrupts Def2’s native conformation (Fig. S2E-F)^[40]^, potentially destabilizing analogous disulfide bridges essential for its function. In contrast, Def1’s structural integrity remains unaltered under identical modifications, consistent with its superior tethering-dependent activity. This divergence suggests that the comparable antimicrobial potency of secreted and tethered Def2 (s-Def2 vs. t-Def2) arises from misfolding or misfunction caused by the appended sequences. To generalize GPI-anchoring as a strategy for AMP optimization, future studies should refine linker flexibility and GPI-anchoring motifs to preserve the structural fidelity of diverse defensins, ensuring membrane localization enhances rather than compromises their antimicrobial mechanisms.

The compromised activity of t-Def2 likely stems from structural perturbations induced by the current linker-GPI fusion (Fig. S2E). Based on our AlphaFold predictions, we propose that shorter (5-8 aa), glycine-rich linkers (e.g., GGGGS or GGSGGS) would: reduce steric interference with defensin folding, maintain proper disulfide bond formation, while preserving membrane anchoring capability. While experimental validation remains for future studies, this structural insight provides a rational framework for optimizing tethering strategies across diverse AMPs. Future work will systematically test linker designs informed by our AlphaFold predictions, combining computational modeling with functional assays to establish general principles for AMP tethering.

The GPI-anchoring strategy demonstrates broad versatility beyond the honeybee defensins central to this work, effectively enhancing the functional display of structurally and functionally diverse peptides. This generalizability is well-supported by complementary studies: our recent systematic evaluation of honeybee venom AMPs established that GPI-tethering consistently boosts immunomodulatory efficacy in *Drosophila* for distinct peptide classes—including not only defensins but also the compact helical apamin and pore-forming melittin ^[41]^. The approach further extends to non-natural scaffolds, as evidenced by GPI-anchored display of the AI-engineered synthetic peptide PAN4, which significantly improved antifungal protection through stabilized cell-surface localization ^[42]^. Perhaps most compellingly, GPI-anchoring proves effective beyond antimicrobial contexts; pioneering work on *Drosophila* neuropeptides demonstrated that tethering signaling molecules like Pigment-Dispersing Factor (PDF) to membranes robustly modulates complex behaviors such as circadian rhythms ^[43,44]^. Collectively, these findings—spanning natural AMPs (defensins, apamin, melittin), synthetic peptides (PAN4), and neuropeptides (PDF) with divergent structures, origins, and functions—solidify GPI-anchoring as a universal platform for concentrating peptides at cell surfaces to amplify their bioactivity.

Looking ahead, refining linker design represents a promising avenue for optimization. The spacer connecting functional peptides to the GPI-anchor may critically influence stability, solvent accessibility, and conformational freedom. Systematic exploration of linker variables—length, flexibility (e.g., glycine-serine repeats), rigidity, and protease resistance—could maximize peptide presentation efficiency and further unlock the platform’s therapeutic potential.

The evolutionary relationship between honeybees and the fruit fly, *Drosophila melanogaster*, indicates a shared biochemical and genetic basis^[45]^. This study offers an opportunity to utilize advanced genetic methods in *Drosophila* to elucidate the genetic relationship between natural defensin-1 and artificially constructed tethered format defensin-1^[15,46]^. The employment of the *Drosophila* model system has significant potential for enhancing our comprehension of the impact of peptide structure on the efficacy of AMPs.

Interestingly, although t-Def1 showed strong biocompatibility under baseline conditions, its expression led to exacerbated intestinal barrier dysfunction under stress paradigms, such as sleep deprivation and DSS challenge, evidenced by increased Smurf phenotype incidence and dye leakage (Fig. 4). This raises the critical question of whether the observed epithelial damage stems from the intrinsic activity of Def1 or is an artifact of the GPI-anchoring strategy itself. While a tethered inert peptide control was not included in the current study, several lines of evidence suggest that the damage is primarily attributable to the membrane-associated bioactivity of Def1 rather than the tethering method per se. First, secreted Def1 (s-Def1), which expresses the same mature peptide without GPI-anchoring, does not induce epithelial dysfunction under identical stress conditions (Fig. 4). Critically, comparative evidence from our prior work argues strongly against a general detrimental effect of membrane tethering. We have consistently expressed diverse GPI-anchored peptides—including the honeybee venom-derived apamin and the synthetic AMP PAN4—under identical experimental paradigms. In these cases, membrane tethering enhanced, rather than compromised, epithelial barrier function without inducing damage ^[41,42]^. This stark phenotypic contrast—robustness versus compromise—demonstrates that epithelial damage is not an inherent outcome of the GPI-anchoring platform, linker design, or the mere presence of a tethered peptide.

Third, the GPI-anchor and linker employed here have been widely used in *Drosophila* without eliciting epithelial toxicity. For example, Rosenbaum et al.^[47]^ and Ruiz et al.^[48]^ demonstrated that GPI-anchored proteins such as chaoptin and grasshopper Lazarillo localize to membranes and function normally without disrupting host physiology. Therefore, the epithelial compromise observed in t-Def1 flies likely results from the sustained membrane localization of a pore-forming AMP, which under stress conditions may increase susceptibility to epithelial damage. This context-dependent trade-off underscores the need to carefully evaluate not only AMP potency but also its subcellular localization and host tolerance, particularly under non-homeostatic conditions. These insights advance our understanding of how membrane anchoring can amplify AMP activity while simultaneously revealing possible vulnerabilities, guiding future AMP engineering toward safer and more targeted applications.

## FIGURE LEGEND

**Fig. S1:** The *Def-1* expression level, and locomotion, courtship index, copulation latency for male s-Def1 and t-Def1 flies. **A,** qRT-PCR Analysis of Def-1 mRNA Expression Levels. Quantitative real-time polymerase chain reaction (qRT-PCR) was performed to measure the relative expression levels of Def-1 mRNA in various genetic backgrounds. The expression levels are normalized to a reference gene (GAPDH) and presented as fold changes. The are data represented as mean ± SEM (n = 3 independent biological replicates). Statistical significance was determined using one-way ANOVA followed by Tukey’s multiple comparisons test. ns, not significant; *****, p < 0.0001. **B,** Climbing Assay. The time taken for climbing to occur was measured in different genotypes of *Drosophila melanogaster*. The data are represented as mean ± SEM (n = 50 flies per genotype). Statistical significance was determined using one-way ANOVA followed by Tukey’s multiple comparisons test. ns, not significant. **C, c**ourtship index measurement. The courtship index, defined as the percentage of time spent in courtship behaviors over a 10-minute observation period, was assessed for various genotypes. The data are represented as mean ± SEM (n = 50 flies per genotype). Statistical significance was determined using one-way ANOVA followed by Tukey’s multiple comparisons test. ns, not significant. CI was measured by [courtship time/ (mating time-courtship time) *100%]. **D,** copulation latency assay. Copulation latency (CL) of male flies expressing Def1, s-Def1, and t-Def1 compared with control (GFP expressing flies) was analyzed using **DrosoMating** and compared with manual recordings. **E,** Pathogen load of female flies expressing t-Def1 compared with control (GFP expressing flies) by *tub-GAL4* following a 12-hour oral feeding of *E.faecal* culture. CFU stands for Colony-Forming Unit. **F,** Pathogen load of female flies +/attP40 compared with Canton-S following a 12-hour oral feeding of *P. aeruginosa* culture. **G,** Standard curve for the c-myc ELISA, generated by simple linear regression (Y = 0.002649*X + 0.2185), where Y represents the Optical Density (OD) at 450 nm and X represents the c-myc concentration in pg/mL. **H,** c-myc protein expression levels, quantified by ELISA, between t-Def1 and t-Def2 expressing female flies driven by *tub-GAL4* ubiquitous expression. Data are presented as individual data points superimposed on a bar plot, with the mean and standard error indicated. **I-J,** Representative confocal microscopy images of the R4 region of the gut from control and t-Def1 expressing flies infected by GFP-tagged *P. aeruginosa* culture. The GFP signal indicates the presence and proliferation of *P. aeruginosa*. DBMs represent the ‘difference between means’ for the evaluation of estimation statistics. Asterisks represent significant differences, as revealed by the Student’s t test (* p<0.05, ** p<0.01, *** p<0.001).

**Fig. S2.** Schematic Representation of Defensin-1 and Defensin-2 Protein Structures and Variants. **A-B,** Structural representations of engineered constructs based on defensin-1 (A, blue) and defensin-2 (B, orange). Individual defensin domains are shown at the top, followed by fusion constructs incorporating a flexible linker (yellow), and ending with constructs containing both the linker and a GPI-anchor (red). N– and C-termini are labeled. The right panels showed the confidences score (pIDDT) of the modeled structures. The very high (pIDDT >90), confident (90 > pIDDT> 70), low (70 > pIDDT > 50), and very low (pIDDT <50) were colored red, light red, light blue, and blue, respectively. **C,** Superposition of full-length defensin-1 fusion proteins with linker and GPI-anchor, highlighting structural conservation. **D,** Structures of defensin-1 and its variants (def-1, sdef-1, tdef-1) with the cysteine residues at the potential disulfide bonds, showing minimal conformational differences. **E,** Structural superimposition of defensin-2 fusion constructs with linker and GPI-anchor, illustrating domain arrangement and overall fold. **F,** Comparison of defensin-2 variants (def-2, sdef-2, tdef-2), showing conserved core structures across constructs.

**Fig. S3:** Representative trajectory diagram of grouped s-Def1 and t-Def1 flies. **A**, Trajectory routes of grouped flies. Representative trajectories of flies expressing s-Defensin-1 (s-Def1) and t-Defensin-1 (t-Def1) are shown. Each color represents the movement path of a single fly within an experimental chamber. Each chamber contains 5 flies. The trajectories were recorded over a defined period to analyze locomotor activity and spatial distribution. The figure illustrates the movement patterns of flies expressing different variants of Defensin-1, highlighting potential differences in locomotor behavior.

**Fig. S4:** Quantification of sleep latency of flies with ubiquitous expression of s-Def1 and t-Def1. **A-B**, Quantification of sleep latency for s-Def1 and t-Def1 expressing flies. **A,** Day time sleep. **B,** Night time sleep. (n=6, n=11 respectively).

**Fig. S5:** Smurf results of female s-Def1 and t-Def1 flies induced by sleep deprivation. **A,** The mean intensity of Smurf assay flies expressing s-Def1 compared with t-Def1 after 24 hours sleep deprivation and 24 hours dye food feeding. (n=7 and 8 respectively)

## METHODS

### Fly stocks and husbandry

*Drosophila melanogaster* was raised on cornmeal-yeast medium at similar densities to yield adults with similar body sizes. The recipe for this medium is as follows: water add up to 5 L, agar 47g, inactive yeast 65.5g, corn flour 232.5g, soy flour 30g, molasses 350 ml, tegosept sol. 35g, propionic acid 12.5ml, phosphoric acid 2.5ml. Flies were kept in 12 h light: 12 h dark cycles (LD) at 25℃ (Zeitgeber Time, ZT 0 is the beginning of the light phase, ZT12 beginning of the dark phase). Flies were flipped every 3-5 days to prevent overcrowding and maintain optimal culture conditions. All mutants and transgenic lines used here and their sources were as follows: *;;tub-GAL4/TM3* (This line was kindly provided by Dr. Lihua Jin, Northeast Forestry University, Harbin, China), *UAS-sDefensin1, UAS-tDefensin1*.

### Bacteria culture

For both normal *P. aeruginosa* (ATCC 27853) and GFP-tagged bacteria culture, 10 mL of Luria-Bertani (LB) broth was inoculated with 100 µL of a frozen bacterial stock at 37 °C. The main procedure was modified based on previous study ^[49]^. For *E. faecalis* (ATCC 29212), MRS medium (LABLEAD, 02-293) was used with the same procedure. The subculture was shaken at 150 rpm overnight and then transferred to a 1 L conical flask for a further overnight incubation. Equal volumes of the subculture were distributed into 500 mL centrifuge tubes and centrifuged at 2,500 × g for 15 min at 4 °C to pellet the bacteria. The supernatant was removed, and the bacterial pellet was resuspended in 5% sucrose water solution. The OD_600_ was measured and adjusted to the desired pathogen load (OD_600_ = 25 in this study).

### Quantitative RT-PCR

The expression levels of *Def* in flies under different genotypes were analyzed by quantitative real-time RT-PCR with SYBR Green qPCR MasterMix kit (Selleckchem). The primers of RT-PCR are, F:5’-GTTCTTCGTTCTCGTGG –3’; R: 5’-CTTTGAACCCCTTGGC –3’. qPCR reactions were performed in triplicate, and the specificity of each reaction was evaluated by dissociation curve analysis. Each experiment was replicated three times. PCR results were recorded as threshold cycle numbers (Ct). The fold change in the target gene expression, normalized to the expression of internal control gene (GAPDH) and relative to the expression at time point 0, was calculated using the 2 ^−ΔΔCT^ method as previously described^[50]^. The results are presented as the mean ± SD of three independent experiments.

### Bacterial infection assay

The main procedure was modified based on previous study and described as follows^[49]^. Flies were flipped every 2-3 days to prevent overcrowding and maintain optimal culture conditions. Flies were starved for 4 h before exposure to bacteria by transferring the flies to empty vials. Place a disc of filter paper on top of food and pipette 100 µL of bacterial culture directly onto the filter disc. For control infections in this study, bacterial culture was replaced with Phosphate-Buffered Saline (PBS). 5 flies were transferred to the sample tube and leave for 12 h infection exposure. To confirm oral infection, first surface-sterilize the flies immediately after bacterial exposure, by placing them in 100 µL of 70% ethanol for 20–30 s. Remove the ethanol and add 100 µL of triple distilled water for 20–30 s before removing the water. Add 100 µL of 1x PBS and homogenize the fly. Transfer the homogenate to the top row of a 96-well plate and add 90 µL of 1x PBS to every well below. Serially dilute this sample to distinguish a range of CFU values. Take 10 µL of the homogenate in the top well and add this to the well below. Repeat this step with the second well, transferring 10 µL to the third well, and so on, for as many serial dilutions as required. Plate the serial dilutions on an LB nutrient agar plate in 2 µL droplets, to ensure all droplets remain discrete. Incubate the LB Agar plates overnight at 37 °C and count visible CFUs. Calculate the number of CFUs per fly by counting the number of colonies present at the serial dilution where 0–40 CFUs are clearly visible. Then check the colony numbers in 10^-5^.

### Honeybee AMP peptides generation

To generate the *UAS-Def1*, *UAS-s-Def1* and *UAS-t-Def1* in this study, peptide cDNAs were chemically synthesized with optimal *Drosophila* codon usage and with an optimal *Drosophila* Kozak translation initiation site upstream of the start methionine (CAAA)^[51]^. Encoded peptides are as follows: *Def1*, MKIYFIVGLLFMAMVAIMAAPVEDEFEPLEHFENEERADRHRRVTCDLLSFKGQVNDSACAANCLSLGKAGGHCEKGVCICRKTSFKDLWDKRFG; *s-Def1*, MSALLILALVGAAVAVTCDLLSFKGQVNDSACAANCLSLGKAGGHCEKGVCICRKTSFKDLWDKRFGGNEQKLISEEDLGN; *t-Def1*, MSALLILALVGAAVAVTCDLLSFKGQVNDSACAANCLSLGKAGGHCEKGVCICRKTSFKDLWDKRFGGNEQKLISEEDLGNGAGFATPVTLALVPALLATFWS LL. These cDNAs were cloned into the pUAS-attB vector; For generation of transgenic *Drosophila*, Vectors was injected into the embryos of flies. The genetic construct was inserted into the attp40 site on chromosome II to generate transgenic flies using established techniques, a service conducted by Qidong Fungene Biotechnology Co., Ltd. (http://www.fungene.tech/).

### Courtship Assays for Copulation Latency (CL) and Courtship Index (CI)

Courtship assay was performed as previously described^[52]^ under normal light conditions in circular courtship arenas 11 mm in diameter, from noon to 4 pm. We utilized the time at which copulation (mating) was initiated to quantify copulation latency (CL). Courtship latency is the time between female introduction and the first obvious male courtship behavior, such as orientation coupled with wing extensions. Once courtship began, courtship index was calculated as the fraction of time a male spent in any courtship-related activity during a 10 min period or until mating occurred. Upon the onset of courtship behavior, the courtship index (CI) was computed as the proportion of time a male dedicated to courtship-related behaviors within a 10-minute observation window or until the initiation of mating. The initiation of mating is defined as the moment at which male flies successfully achieve mounting on females.

### Single fly locomotion assay

In the single locomotion experiment, 20 female flies aged 5-7 days were individually placed into a plate with single chambers, and a 30-minute video was recorded to track the movement of each fly. To detect and quantify the activity of flies, we have developed the Fly Trajectory Dynamics Tracking (FlyTrDT) software. This is a custom-written Python program that utilizes the free and open-source OpenCV machine vision library and Python Qt library. The FlyTrDT software simultaneously records the trajectory information of each fly and calculates various indicators of the group at a certain period. For each frame acquired, the moving fly is segmented using the binarization function from the OpenCV library. Subsequently, a Gaussian blur and morphological closing and opening operations were performed on the extracted foreground pixels to consolidate detected features and reducing false positives and negatives. Finally, the extraction of fly outlines was achieved using the contour detection algorithm in the OpenCV library.

### Group fly locomotion assay

We adapted the fly Group Activity Monitor (flyGrAM) system described by Scaplen et al. (2019)^[53]^ to develop Fly Trajectory Dynamics Tracking (FlyTrDT), a customized software for quantifying group locomotion behavior. For experiments, 27 female *Drosophila melanogaster* (5–7 days post-eclosion) were divided into three cohorts (9 flies/group) and transferred to separate circular behavioral chambers (30 mm diameter, 2 mm height), constructed from transparent acrylic. Flies were reared at 18°C prior to eclosion and maintained at 25°C post-eclosion for behavioral assessments. For experimental setup and imaging preparation, a custom LED backlight array beneath the chambers provided uniform illumination through a 2-mm transparent acrylic diffuser. Fly movement was recorded at 30 Hz (1920 × 1080 resolution) using a mounted camera. Following a 30-minute acclimatization period to exclude individuals with motor defects, 30-minute videos were acquired for analysis. FlyTrDT, an open-source Python-based program integrating OpenCV and Qt libraries, was executed on a Windows workstation (minimum 8 GB RAM, dual-core 2.1 GHz processor). For each frame, foreground pixels corresponding to flies were isolated via OpenCV’s binarization function. Gaussian blur and morphological operations (closing/opening) enhanced feature detection while minimizing noise. Fly contours were identified using OpenCV’s contour detection algorithm. Trajectories were reconstructed by matching current contours to prior positional, directional, and velocity data, with predictive modeling to resolve occlusions.

### Climbing assay

For climbing assay, we modified the conventional RING assay and designed conventional Fast Inexpensive Climbing Test (FICT)^[54,55]^. In brief, approximately forty 20-day-aged flies were placed in an empty vial and were tapped to the bottom of the tube. After tapping of flies, we recorded 10 seconds of video clip. This experiment was done five times with 1-minute intervals. With recorded video files, we captured the position of flies 10 seconds after tapping the vial. This captured image file was then loaded in ImageJ to perform particle analysis. For quantifying the location of flies inside a vial, we used the “analyze particles” function of ImageJ^[55,56]^. The position of pixels was normalized by height of vial then only the particles above the midline (4 cm) of vial were counted.

### Single-fly sleep and circadian rhythm recording

96-well white Microfluor 2 plates (Fishier) with 400 μl of food (5% sucrose and 1% agar) were loaded with adult male flies (aged 3–5 days). Flies were entrained to the 12 h:12 h LD cycles for four days at 25 °C to record sleep behavior, then changed to constant darkness for 5-6 days to record circadian rhythms in the absence of light inputs. The fly movement was monitored using a camera at 10s intervals, and the data were then used by the sleep and circadian analysis program SCAMP to analyze sleep and circadian rhythm^[57–59]^. It calculates activity by shifting the position of *Drosophila* every 10 seconds and calculates sleep using the standard definition (*Drosophila* is recorded as asleep if it remains motionless for at least 5 minutes).

### Smurf assay

For the Smurf assay, we used 3% (v/v) Food Blue No.1 aluminum lake (Aladdin, Cat# F336821). The blue dye was mixed directly into standard fly food, and flies were fed for 6 or 20 hours depending on the specific experimental condition (with or without prior stress). Flies were imaged using an Olympus SZ61 stereomicroscope under uniform lighting. Images were captured at a fixed magnification. Representative Smurf images are shown in Figure 4^[60]^. For flies under sleep deprivation, a vortex machine (CHANGZHOU ENPEI INSTRUMENT MANUFACTURING CO., LTD., NY-5SX) was applied with a routine of 2-second 1500 rpm vortex following a one-minute rest^[61]^, flies were transferred in tubes containing normal food and went through a 24-hour sleep deprivation. Dextran sulfate sodium (DSS; Coolaber, Cat# 9011-18-1) was dissolved with blue dye food to a final concentration of 3% (w/v) and mixed thoroughly into fly food prior to administration. Flies were fed this DSS-containing food for 24 hours before being assessed for gut permeability^[62]^.

### Structural modeling of engineered Def-1 and Def-2

Genetic Constructs and Insertion Loci: Detailed maps of all AMP constructs (Def1, s-Def1, t-Def1) including signal peptides, linker sequences, and GPI-anchor regions have been included as Figure 1. All constructs were inserted into the well-characterized attP40 site on chromosome II using PhiC31-mediated integration to ensure consistent expression and reduce positional effects. The sequences relevant to native Def-1, sDef-1, tDef-1, native Def-2, sDef-2, and t-Def2 were N-VTCDLLSFKGQVNDSACAANCLSLGKAGGHCEKGVCICRKTSFKDLWDKRF G-C, N-VTCDLLSFKGQVNDSACAANCLSLGKAGGHCEKGVCICRKTSFKDLWDKRFGGNEQKLISEEDLGN-C, N-VTCDLLSFKGQVNDSACAANCLSLGKAGGHCEKGVCICRKTSFKDLWDKRFGGNEQKLISEEDLGNGAGFATPVTLALVPALLATFWSLL-C, N-VTCDVLSWQSKWLSINHSACAIRCLAQRRKGGSCRNGVCICRK-C, N-VTCDVLSWQSKWLSINHSACAIRCLAQRRKGGSCRNGVCICRKGNEQKLISE EDLGN-C, N-VTCDVLSWQSKWLSINHSACAIRCLAQRRKGGSCRNGVCICRKGNEQKLISEEDLGNGAGFATPVTLALVPALLATFWSLL-C were used for the prediction of overall structure by using AlphaFold3 in AlphaFold web server (https://alphafoldserver.com/)^[63]^. The first ranked models were then structurally analyzed. All structural figures were generated by using chimeraX^[64]^.

### Immunostaining

5 days after eclosion, the *Drosophila* was infected as described previously and the gut was taken from adult flies and fixed in 4% formaldehyde at room temperature for 30 minutes. The sample was than washed three times (5 minutes each) in 1% PBT and then blocked in 5% normal goat serum for 30 minutes. Subsequently, the sample was incubated overnight at 4℃ with primary antibodies in 1% PBT, followed by the addition of fluorophore-conjugated secondary antibodies for one hour at room temperature. For DAPI staining, the gut was incubated in DAPI containing 1%PBT for 10 minutes at room temperature, followed by three times wash. Finally, the gut was mounted on plates with an antifade mounting solution (Solarbio) for imaging purposes. Samples were imaged with Zeiss LSM880. Antibodies were used at the following dilutions: Chicken anti-GFP (1:500, Invitrogen), mouse anti-c-myc (1:200, Abcam), Alexa-488 donkey anti-chicken (1:200, Jackson ImmunoResearch), Alexa-647 donkey anti-mouse (1:200, Invitrogen), DAPI (1:1000, Invitrogen).

### c-myc ELISA Quantification

c-myc protein expression levels were quantified using the Fruit Fly c-myc ELISA Kit (LUPUKEJI). For sample preparation, 25 flies from each genotype were homogenized in 400 µL of saline solution. The homogenates were then centrifuged at 3000 x g for 10 minutes, and the resulting supernatant was used for the ELISA. Optical density (OD) was measured at 450 nm, and c-myc concentrations were calculated according to the manufacturer’s instructions.

### Statistics

All analysis was done in GraphPad (Prism9). For two group comparing, we used unpaired two-tailed Student’s *t*-tests, and for multiple group comparisons, one-way ANOVA with Tukey’s post hoc test was applied. Survival curves were analyzed using the log-rank (Mantel–Cox) test. In addition to conventional null hypothesis significance testing (NHST)^[65]^, we applied the estimation statistics framework to all two-group comparison datasets, including bacterial infection assays, immunostaining, and Smurf assay results. We used the estimation graphics platform (https://www.estimationstats.com) to calculate effect sizes, including mean differences (DBM) and 95% confidence intervals, and to visualize the full distribution of observed values and effect precision. This method complements NHST by shifting the focus toward effect size and data distribution rather than binary significance thresholds. All data are reported as mean ± SEM, and exact *n* values and *p*-values are indicated in figure legends. In climbing assays, mean values were compared by one-way ANOVA, each figure shows the mean ± standard error (SEM) (\**\*\*\* = p<0.0001, *** = p < 0.001, ** = p < 0.01, * = p < 0.05*, n.s. stands for non-significant differences). Besides traditional *t*-test for statistical analysis, we added estimation statistics for all two group comparing graphs. In short, ‘estimation statistics’ is a simple framework that—while avoiding the pitfalls of significance testing—uses familiar statistical concepts: means, mean differences, and error bars.

More importantly, it focuses on the effect size of one’s experiment/intervention, as opposed to significance testing ^[66]^. Thus, we conducted a reanalysis of all our two group data sets using both standard *t* tests and estimate statistics. In 2019, the Society for Neuroscience journal eNeuro instituted a policy recommending the use of estimation graphics as the preferred method for data presentation ^[67]^.

### Data and code availability

● All data reported in this paper will be shared by the lead contact upon request.
● This paper does not report original code. The URL of the codes used in this paper are listed in the key resources table.
● Any additional information required to reanalyze the data in this paper is available from the lead contact upon request.

## ACKNOWLEDGEMENT

We are very appreciative to the colleagues who supplied us with several fly strains; especially Dr. Lihua Jin (Northeast Forestry University, Harbin, China). The GFP-tagged *P. aeruginosa* was kindly given by Dr. Dominique Ferrandon. Stocks obtained from the Bloomington Drosophila Stock Center (NIH P40OD018537) were used in this study. The fly stock was obtained from Korea Drosophila Resource Center and National Drosophila Resource Center of China. This work was supported by Startup funds from HIT Center for Life Science to WJK.

## AUTHOR CONTRIBUTIONS

**Conceptualization**: Woo Jae Kim.

**Data curation**: Yanan Wei, Yanying Sun, Xinyue Zhou, Doyoun Kim, Woo Jae Kim.

**Formal analysis**: Yanan Wei, Yanying Sun, Doyoun Kim, Woo Jae Kim.

**Funding acquisition**: Woo Jae Kim.

**Investigation**: Woo Jae Kim.

**Methodology**: Jihyeon Lee, Jeong Kyu Bang, Woo Jae Kim.

**Project administration**: Woo Jae Kim.

**Resources**: Jihyeon Lee, Jeong Kyu Bang, Woo Jae Kim.

**Supervision**: Woo Jae Kim.

**Validation**: Yanan Wei, Yanying Sun, Xinyue Zhou, Doyoun Kim, Woo Jae Kim.

**Visualization**: Yanan Wei, Yanying Sun, Xinyue Zhou, Doyoun Kim, Woo Jae Kim.

**Writing – original draft**: Yanan Wei, Doyoun Kim, Woo Jae Kim.

**Writing – review & editing**: Yanan Wei, Yanying Sun, Woo Jae Kim.

