## Supplementary figures and images for "Membrane Tethering of Honeybee Antimicrobial Peptides in *Drosophila* Enhances Pathogen Defense at the Cost of Stress-Induced Host Vulnerability"

### Supplemental Figure 1

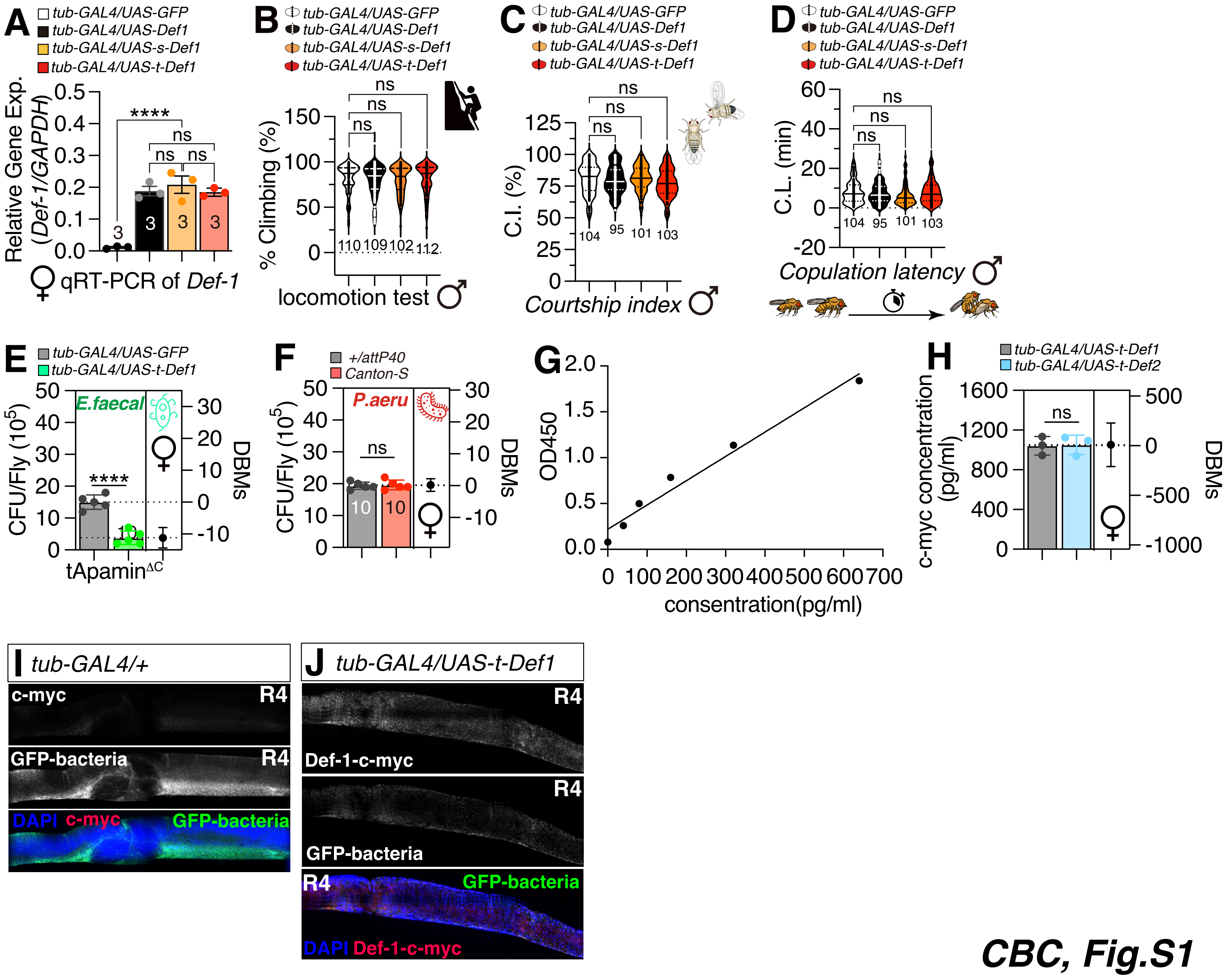

### Supplemental Figure 2

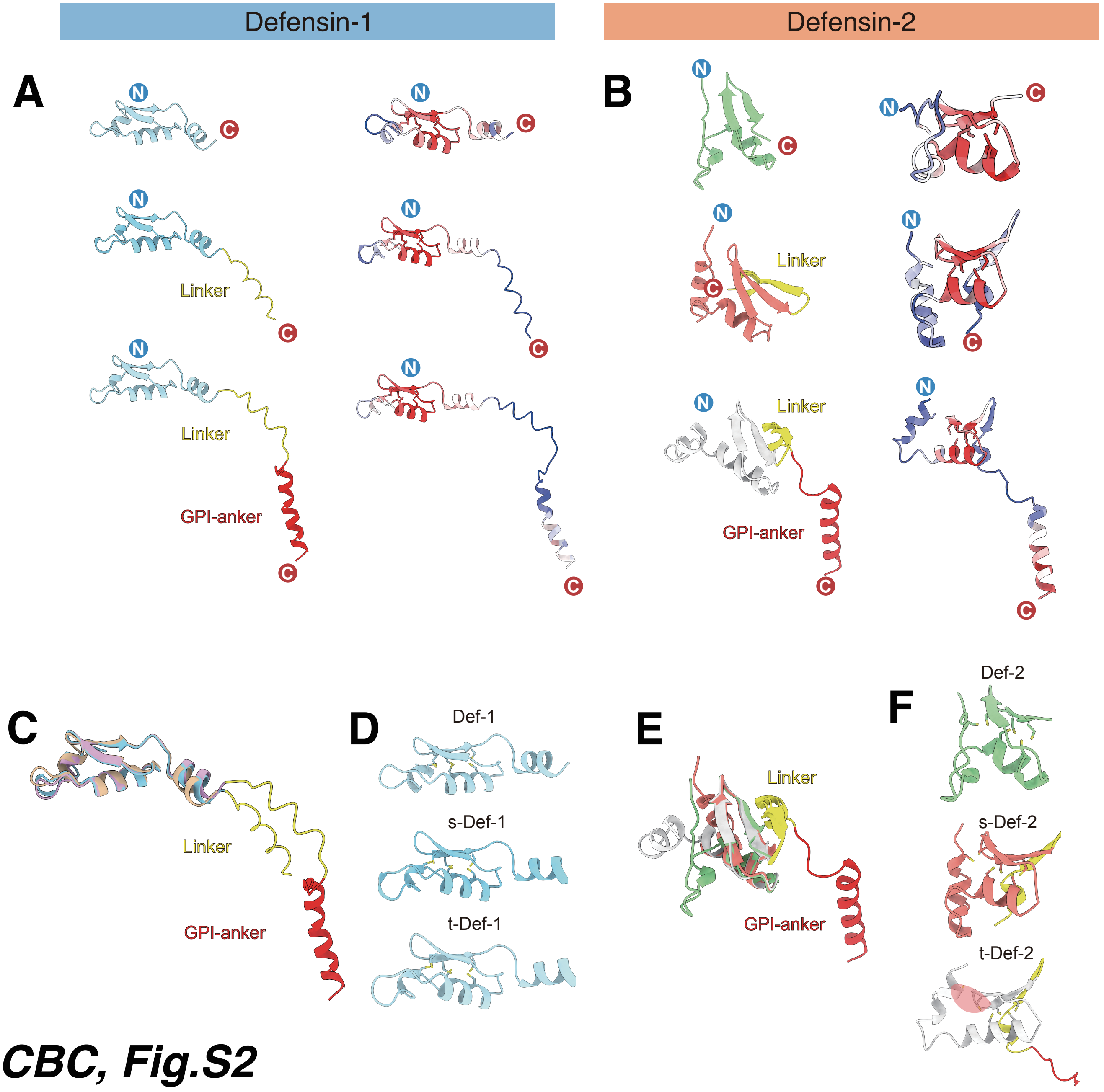

### Supplemental Figure 3

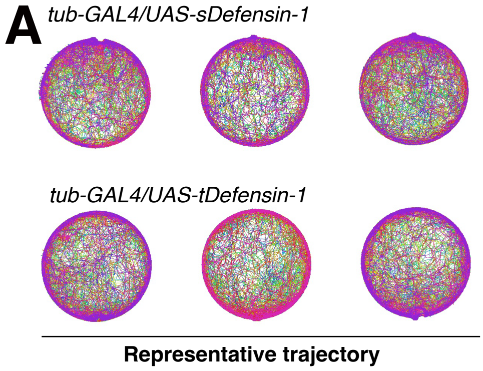

### Supplemental Figure 4

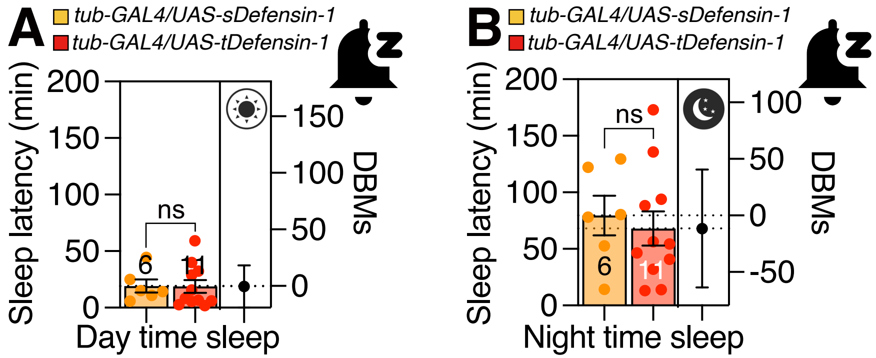

### Supplemental Figure 5

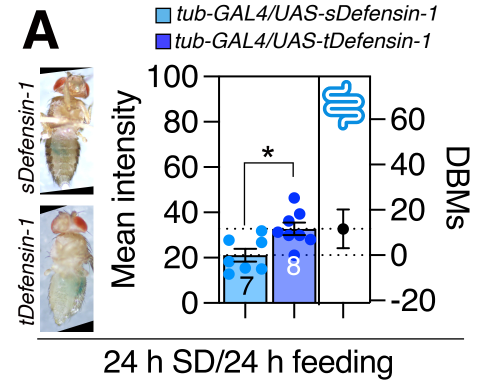
