## Supplemental Table 1 for "Membrane Tethering of Honeybee Antimicrobial Peptides in *Drosophila* Enhances Pathogen Defense at the Cost of Stress-Induced Host Vulnerability"

| Construct | Type |
| --- | --- |
| UAS-Def1 | Nucleotide |
| | Peptide |
| | Features |
| UAS-s-Def1 | Nucleotide |
| | Peptide |
| | Features |
| UAS-t-Def1 | Nucleotide |
| | Peptide |
| | Features |

**Sequence (5'→3' or N→C)**

ATGAAAATTTATTTTATTGTTGGTTTATTGTTTATGGCGATGGTGGCTGCTC
CGGTAGAAGACGAGTTTGAGCCGCTGGAGCACTTTGAGAACGAGGAGCGT
GCCGACCGCCACCGCCGTGTCACCTGCGACCTGCTGTCTTTCAAGGGCCAA
GTGAACGACTCGGCCTGCGCCGCGAACTGCCTGTCGCTGGGCAAGGCTGG
TGGCCACTGCGAGAAGGGTGTGTGCATCTGCCGCAAGACGTCTTTCAAGG
ACCTGTGGGACAAGCGGTTTGGTGA

MKIYFIVGLLFMAMVAIMAAPVEDEFEPLEHFENEERADRHRRVTCDLLSFKG
QVNDSACAANCLSLGKAGGHCEKGVCICRKTSFKDLWDKRFG

Native defensin1 with endogenous signal peptide

ATGTCGGCGCTGCTGATCCTGGCCCTGGTGGGAGCCGCCGTCGCCGTCACC
TGCGACCTGCTGTCTTTCAAGGGCCAAGTGAACGACTCGGCCTGCGCCGCG
AACTGCCTGTCGCTGGGCAAGGCTGGTGGCCACTGCGAGAAGGGTGTGTG
CATCTGCCGCAAGACGTCTTTCAAGGACCTGTGGGACAAGCGGTTTGGTGG
TGGCGAGAAGCAGAAGCTGATCTCGGAGGAGGACCTGGGTAAC

MSALLILALVGAAVAVTCDLLSFKGQVNDSACAANCLSLGKAGGHCEKGVCI
CRKTSFKDLWDKRFGGNEQKLISEEDLGN

α-Bgtx signal peptide + Def1 + hydrophilic linker

ATGTCGGCGCTGCTGATCCTGGCCCTGGTGGGAGCCGCCGTCGCCGTCACC
TGCGACCTGCTGTCTTTCAAGGGCCAAGTGAACGACTCGGCCTGCGCCGCG
AACTGCCTGTCGCTGGGCAAGGCTGGTGGCCACTGCGAGAAGGGTGTGTG
CATCTGCCGCAAGACGTCTTTCAAGGACCTGTGGGACAAGCGGTTTGGTGG
TGGCGAGAAGCAGAAGCTGATCTCGGAGGAGGACCTGGGTAACGGTGCCG
GCTTCGCCACCCCGGTCACCCTGGCCCTGGTCCCGGCGCTGCTGGCGACCT
TCTGGTCGCTGCTGTGA

MSALLILALVGAAVAVTCDLLSFKGQVNDSACAANCLSLGKAGGHCEKGVCI
CRKTSFKDLWDKRFGGNEQKLISEEDLGNGAGFATPVTLALVPALLATFWSLL

s-Def1 sequence + GPI anchor
